# Establishment of a Human RNA Pol II Pausing System and the Identification of O-GlcNAc Cycling Regulating Pol II Pausing and Elongation

**DOI:** 10.1101/2020.01.23.917237

**Authors:** Brian A. Lewis, David Levens

## Abstract

Paused RNA polymerase II is the major regulated step of transcription in metazoans. We describe here a unique human cell-free transcription system that recapitulates RNA pol II pausing and assemble paused pol IIs on the human CMV IE, SV40, and heat shock promoters, all is the case in vivo. We then use the system to show that PARP-1 and CDK12/13 inhibitors directly affect pausing and elongation. We then show that O-GlcNAcylation is required for the establishment of a paused pol II: inhibition of OGT allowed pol II to bypass pausing and begin to elongate. Addback of rOGT or the pausing factor NELF re-established the pausing. In vivo nascent RNA measurements showed that OGA inhibition blocks elongation. These data show that cell-free systems can recapitulate RNA pol II pausing, that PARP-1 and CDK12/13 directly regulate RNA pol II elongation, and identify O-GlcNAc cycling regulating both pausing and pause release.

## Introduction

Our understanding of transcriptional regulation by RNA polymerase II has undergone major revisions over the past several years. The predominant view of pol II transcriptional regulation assigned the primary regulatory point to the promoter region of a gene (consisting of various core promoter elements and activator binding sites) and the congregation of general transcription factors that were then recruited by an activator. The ordered assembly of factors at the promoter forms the preinitiation complex (PIC). This view in part arose from the identification of the general factors and the eventual reconstitution of pol II-dependent transcription. Consequently, the formation of the PIC was thought to be the major regulatory step at promoters: upon initiation of transcription, pol II immediately enters productive elongation.

Over the past 10 years genome-wide studies have shown that many, if not the majority of promoters, from *Drosophila* to humans, are regulated at the paused pol II step, where an engaged pol II sits between +30 and +75 downstream of the transcriptional start site (Adelman and Lis 2012). Thus, it is important to understand the nature and regulation of the paused pol II both as a fundamental problem and because many diseases are now considered “diseases of uncontrolled transcriptional elongation” (Chen et al. 2018).

A paused pol II is thought to be established by two factors, DSIF and NELF (Wada et al. 1998b, 1998a; Yamaguchi et al. 1999), that together bind to pol II and effectively misalign the active site of pol II from the DNA template preventing nucleotide incorporation and elongation (Bernecky et al. 2017; Vos et al. 2018). Pol II release occurs via the phosphorylation of DSIF and NELF by P-TEFb (Yamada et al. 2006; Fujinaga et al. 2004), after which NELF is lost, and the polymerase moves into a productive elongation state (for review see Guo and Price 2013).

However, this model is incomplete. The addition of purified DSIF and NELF does not establish a paused pol II, but only slows elongation (Cheng and Price 2007; Renner et al. 2001). These data imply that additional factors are required to establish a paused pol II. Secondly, another catalytic activity, poly-ADP ribose polymerase 1 (PARP-1), also regulated pol II release in cells (Gibson et al. 2016), and so elongation is no longer simply due to P-TEFb activity.

More recently, an O-GlcNAcylation requirement in transcription has emerged. O-GlcNAc is found on serine and threonine resides, often in a mutually exclusive relationship with phosphorylation. There is only one transferase, the O-GlcNAc transferase (OGT) and one removal enzyme, O-GlcNAc aminidase (OGA) (Bond and Hanover 2013). High densities of OGA, OGT and O-GlcNAc co-localize with paused promoters in humans, *M. musculus, Drosophila*, and *C. elegans* (Resto et al. 2016; Love et al. 2010; Deplus et al. 2013; Krause et al. 2018). Previous experiments suggested that OGA activity is necessary for pol II elongation in a human cell-free system (Resto et al. 2016). For OGA to have a substrate, OGT must have acted prior to OGA, but it was not clear whether OGT activity occurred before or after PIC formation.

A cell-free transcription system that recapitulates pol II pausing would be highly desirable, enabling the dissection of the pausing machinery and the regulation of pause establishment and release. We describe here a human cell-free transcription system (CFS) that contains an actively engaged paused pol II which can be released via P-TEFb or SEC (which contains P-TEFb), Sarkosyl, or excess SPT5. This system was used to identify other, previously unknown components of the pausing machinery. Specifically, inhibition of the catalytic activity of OGT results in the loss of paused pol II, which instead immediately elongates, effectively bypassing the pausing step. This pausing defect can be rescued by the addition of rOGT or purified human NELF complex, suggesting that NELF-dependent pausing requires OGT. The CFS shows that pol II pausing can be recapitulated in human derived nuclear extracts. Additionally, these data show that the decision to pause is an intrinsic part of the initiation process and is predicated on the action of OGT, identifying O-GlcNAc cycling as a necessary regulatory event in RNA pol II pausing and release.

## Results

### The cell-free system recapitulates in vivo elongation events

Recently, one of us developed a nuclear extract from HeLa cells and the elongation properties of this extract were used to identify OGA as an elongation factor (Resto et al. 2016). One interesting observation from those experiments was that the RNA products below the tRNA band at +75 were not affected by the inhibition of either OGA or P-TEFb, whereas full-length elongation was completely blocked. Lastly, the specific localization of these RNAs between +20 and +75 is exactly where one would expect to find a paused pol II population (for review see Kwak et al. 2013). The following experiments set out to determine the extent to which the CFS recapitulated in vivo behaviors and whether these polymerases between +20 and +75 were in fact paused.

### Elongation inhibitors that block elongation in vivo show elongation defects in the CFS

In order to establish the legitimacy of the CFS, it was necessary to determine whether the system recapitulated the effects of known in vivo elongation inhibitors. Several groups have shown in vitro and in vivo that THZ1 inhibitor blocks pol II elongation (Nilson et al. 2015; Kwiatkowski et al. 2014). THZ1 shows the highest specificity for the CDK7 kinase (IC50 = 27nM) while also inhibiting CDK12/13 at higher concentrations (IC50 = 250nM) (Kwiatkowski et al. 2014). Addition of THZ1 (4 μM final) to the CFS showed a clear elongation block of full-length 548 nt RNA product (lane 4, Fig. 1B). Additionally, the RNA population between +20 and +75 decreased while there is an increase in RNAs chased beyond the tRNA band at +75 [Note that tRNAs in the extract are factitiously labeled by CCA tRNA nucleotidyl transferase and the ^32^P-CTP (Adamson et al. 2003)]. The expected elongation block with the CDK9 inhibitor flavopiridol was also observed (lane 3, Fig. 1B) (Cheng et al. 2012).

**Figure 1.**
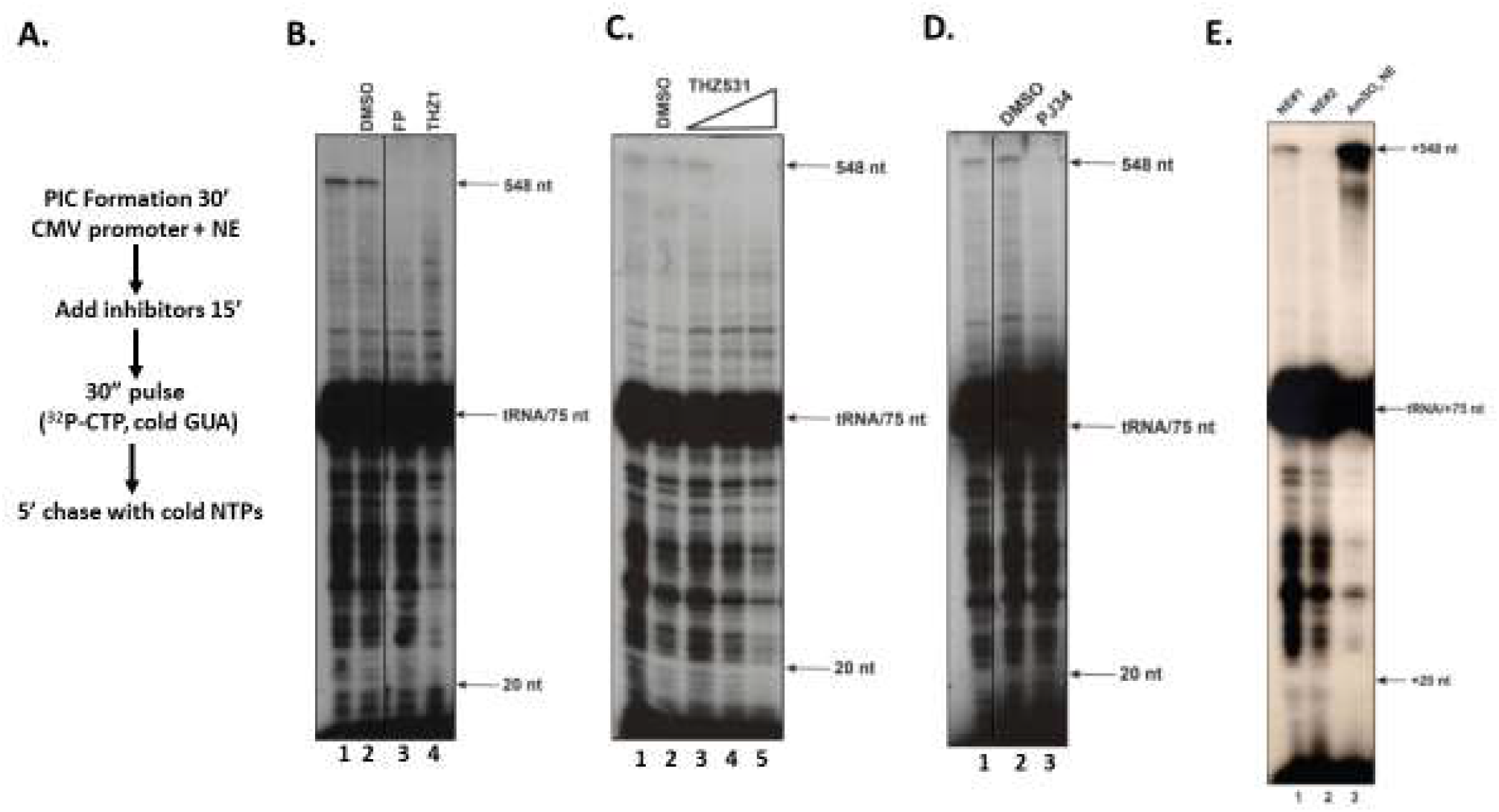
Cell-free system is dependent on CDK7, CDK9, CDK12/13, and PARP-1. A. Schematic of steps in the experiments in panels B-D. B. Shown is a pulse-chase assay using the DMSO solvent alone, 1 μM CDK9 inhibitor flavopiridol (FP), or 4 μM CDK7 inhibitor THZ1 in DMSO. Transcription products were analyzed on an 8% urea/TBE gel followed by autoradiography. Vertical black line between lanes 2 and 3 indicates a deletion of intervening lanes relative to the original autoradiograph. C. Shown is a pulse-chase assay using the DMSO solvent alone and a titration of the CDK12 inhibitor THZ531 (0.4 μM, 4 μM, 40 μM final concentrations). Each THZ531 addition was in the same volume of DMSO as in lane 2. D. Shown is a pulse-chase assay using the DMSO solvent alone or the PARP-1 inhibitor PJ34 (80 μM final concentration). Vertical black line between lanes 1 and 2 indicates a deletion of intervening lanes relative to the original autoradiograph. E. Shown is a pulse-chase assay of two nuclear extracts made as indicated in Materials and Methods (lanes 1 and 2) and compared to an extract made with an ammonium sulfate based protocol (lane 3) (Adamson et al. 2003).

The CFS was assessed for the effects with the CDK12/13 inhibitor THZ531 (IC_50_=158 nM/69 nM for CDK12/13 and IC_50_=8.5 μM/10.5 μM for CDK7 and CDK9, respectively), that also blocked pol II elongation in vivo (Zhang et al. 2016). Titration of increasing concentrations of THZ531 (in the same volume of DMSO) into the CFS showed effects similar to those of THZ1: a reduction of short RNAs between +20 and +75, an increase in RNAs above +75, and a block of full-length RNA synthesis (compare lane 2 to lanes 3-5, Fig. 1C).

Kraus and colleagues have described a positive role for PARP-1 in pol II elongation in vivo (Gibson et al. 2016). To examine the CFS for possible dependence on PARP-1, we added the PARP-1 inhibitor PJ34 to the CFS post-PIC formation, which resulted in a clear block in elongation with loss of the 548 run-off product (lane 3, Fig. 1E), identical to flavopiridol (Fig. 1B), while having no effect on the +20/+75 RNA population; DMSO alone had no effect (lane 2, Fig. 1D).

These experiments show that the CFS can recapitulate known elongation defects previously documented in vivo with flavopiridol, THZ1, THZ531, and PJ34 (Nilson et al. 2015; Zhang et al. 2016; Kwiatkowski et al. 2014; Gibson et al. 2016).

Lastly, we compared our extracts to an extract made with an ammonium sulfate-based protocol and which has previously been used to study RNA pol II elongation (Adamson et al. 2003; Cheng and Price 2007, 2009; Renner et al. 2001). We assayed both types of extracts side-by-side and confirmed that the behaviors of these ammonium sulfate-derived extract (AmSO_4_ NE) is entirely different: elongation is extremely efficient with little to no accumulation of RNAs below +75, as compared to our extracts (Figure 1E, compare lanes 1 and 2 to lane 3). This experiment further suggested that we had developed an extract that was functionally and biochemically distinct.

### Defining a paused pol II population

To further address whether the RNAs between +20 and +75 represent a paused pol II population (delineated by the red brackets in Figure 2), we undertook several litmus tests. If these are paused polymerases, then one would expect that they satisfy several criteria: the paused pol II population should not chase into longer RNAs unless appropriately stimulated, should be sensitive to the detergent Sarkosyl and the addition of DSIF (a known pausing factor), and P-TEFb should stimulate release of the paused pol II and the synthesis of longer RNAs. Additionally, promoters known to establish a paused pol II in vivo should do so in the CFS.

**Figure 2.**
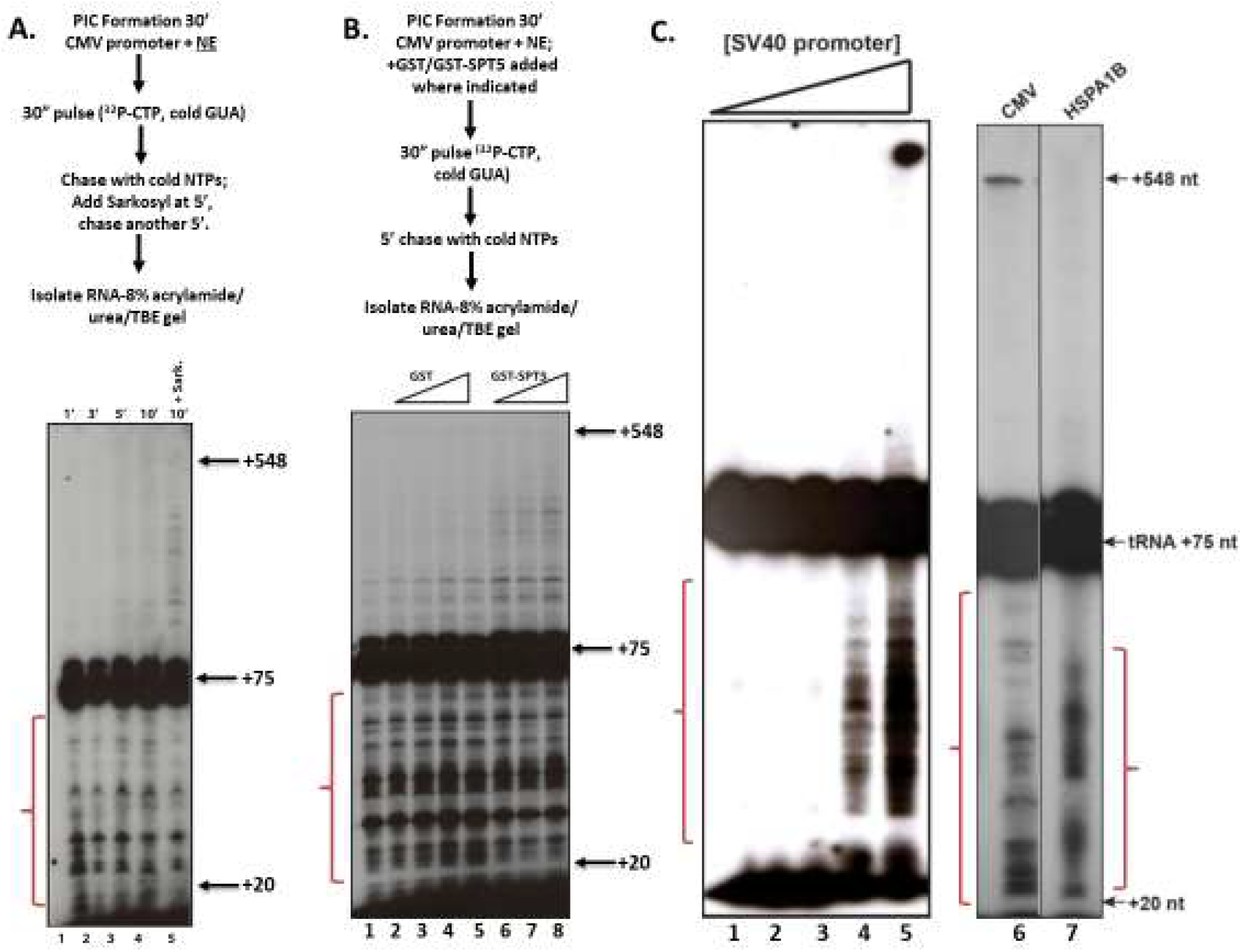
Sarkosyl and excess SPT5 elicit release of paused pol II. A. Pausing system is sensitive to Sarkosyl and paused pol II does not chase. On the left is a schematic of steps in the experiment in this panel. Shown are pulse-chase assays allowed to chase for either 1, 3, 5, or 10 minutes (lane 1-4) or for 5’, followed by the addition of 0.6% final concentration of Sarkosyl, followed by an additional 5’ chase (lane 5). B. Pol II is released by excess SPT5. Shown on the left is a schematic of steps in the pulse-chase assay on the right. Equivalent amounts of rGST or rGST-SPT5 were titrated into PICs that had been previously formed for 30’. After a 15’ incubation, the pulse-chase assay was performed on each titration. C. SV40 and heat shock promoters establish a paused pol II. SV40 promoter DNA (10, 25, 50, 100, 200 ng; lanes 1-5, respectively) was incubated with nuclear extracts for 30’ followed by a standard pulse-chase assay. Lane 6 is a CMV control pulse/chase assay and lane 7 is a pulse/chase assay using the human HSPA1B promoter. These two lanes were merged from the same autoradiograph. Transcription products were analyzed on an 8% urea/TBE gel followed by autoradiography.

### Sarkosyl releases the paused pol II

The release of paused pol II with the detergent Sarkosyl has been documented previously, in both in vivo nuclear run-on assays and in Drosophila nuclear extracts (Rougvie and Lis 1988; Li et al. 2013). The proposed mechanism of Sarkosyl release is that treating the paused pol II with a mild detergent removes the pausing factors thereby permitting pol II release and elongation (Wissink et al. 2019).

We set out to examine the stability of the putative paused pol II population and the effects of Sarkosyl on that population. A pulse-chase time course was done to determine the stability of the paused pol II. A chase with unlabeled CTP was allowed for 1, 3, 5, or 10 minutes (Fig. 2A, lanes 1-4). The +20/+75 RNAs are remarkably stable, with minimal incorporation into longer products even after a 10’ chase. We next added Sarkosyl to the CFS after a 5’ chase followed by an additional 5’chase, which released a fraction of +20/+75 RNAs into longer species, in contrast to a 10’ chase without Sarkosyl (compare lanes 4 and 5, Fig. 2A). Note that because of the excess of cold NTPs in the chase, the longer labeled RNAs here must have originated from the short, pulsed RNA population that elongated after the Sarkosyl addition.

This experiment shows that the +20/+75 RNAs are nascent transcripts engaged with paused polymerases that can elongate after Sarkosyl addition, as described in vivo, but otherwise do not chase even after 10’. This is in contrast to other published elongation systems where the entire pol II population will chase within 3 minutes to full-length RNA products (Cheng and Price 2007).

### GST-SPT5 stimulates elongation in the cell-free system

The DSIF complex, a heterodimer of SPT4 and SPT5, is one of two known pausing factors (Yamaguchi et al. 2013), but can also stimulate elongation (Chen et al. 2009; Yamada et al. 2006). As such, one might expect that the addition of rSPT5 to the CFS would stimulate elongation, either by a squelching mechanism where it titrated out other pausing factors, or through its positive elongation activity. To that end, equivalent amounts of either rGST or rGST-SPT5 were titrated into the CFS after PIC formation. After a 10’ incubation, the standard pulse-chase assay was performed (Fig. 2B). The addition of GST-SPT5 stimulated release of the +20/+75 RNA/paused pol II population, as there is a diminution of RNA around +20 and a concomitant increase of RNAs longer than +75 (Fig. 2B, lanes 6-8), while GST protein alone had little effect (Fig. 2B, lanes 2-5). Although the stimulation of elongation was not efficient as no increase in full-length products was seen, this experiment indicates that the CFS contains a pol II population that elongates in the presence to excess SPT5 as we predicted.

### Pausing occurs in the CFS using a human heat shock promoter and SV40 promoter

We next asked whether other promoters known to harbor a paused pol II would show this behavior in the CFS. The SV40 promoter was one of the first promoters suggested to have a paused pol II (Skolnik-David and Aloni 1983). Titration of the SV40 promoter DNA into the CFS clearly showed the establishment of a paused pol II population between +20 and +75 (lanes 1-5, Fig. 2C).

We extended these observations to the well-known pausing on heat shock promoters. We used the human HSPA1B promoter (Bunch et al. 2014) in our standard pulse/chase assay and found that a discrete paused pol II population existed between +20 and +75 nt, (Fig. 2C, lanes 6-7). These data show that pausing occurs as a default event on three different promoters (CMV, SV40, and HSPA1B) harboring a paused pol II in vivo.

### P-TEFb and Super Elongation Complex stimulate pause release in the CFS

We next determined whether the CFS and the paused pol II population would respond to the addition of recombinant P-TEFb, which is required for pause release in vivo. In order to assay P-TEFb directly, the endogenous activity must first be removed from the CFS. Using a biotinylated-CMV promoter template conjugated to magnetic beads, one can isolate various steps in the transcription processes, wash off soluble factors, and then add back the activity in question to assess its function (Resto et al. 2016; Adamson et al. 2003; Cheng and Price 2009). As indicated in Fig. 3A, a 30” pulse was used to label RNA and transcription was stopped with EDTA. The beads were isolated and washed with a buffer containing 60 mM KCl, and then resuspended in transcription buffer. As shown in lane 1, Fig. 3C, there is no full-length product synthesized after the addition of NTPs for the chase step. The addition of a concentrated 0.3M KCl elongation fraction eluted from a P11 column (P.3c) (Wada et al. 1998a), stimulated synthesis of full-length RNA (lane 2, Fig. 3C). The addition of recombinant P-TEFb, purified from Sf9 cells (Fig. 3B), further stimulated elongation (lane 3, Figure 3C), showing that the paused polymerases isolated on the immobilized templates are active and capable of being released into productive elongation and synthesizing a full-length RNA.

**Figure 3.**
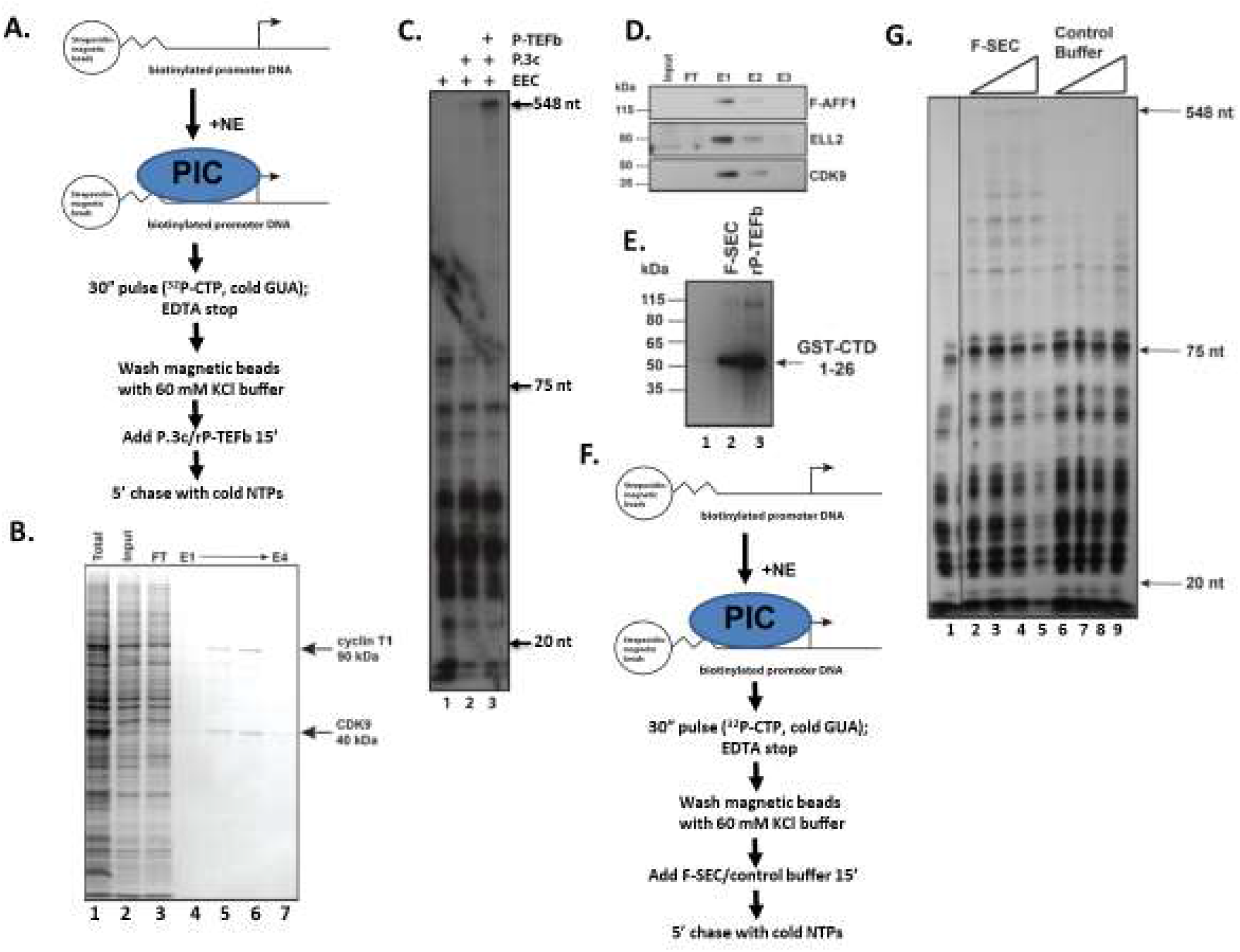
Super Elongation Complex (SEC) and P-TEFb stimulate release of paused RNA pol II. A. Shown is a schematic of steps in the immobilized template assays in panels C and F. B. Purification of rPTEFb. Baculovirus expression vectors containing human P-TEFb and cyclin T1 were expressed in Sf9 cells and purified by Ni-Sepharose affinity chromatography. C. Pol II elongation is stimulated by rP-TEFb. Pulsed immobilized templates were isolated after they had been isolated and washed to remove ^32^P-CTP/GUA and nuclear extract. The indicated factors (P.3c is a concentrated P11 0.3M elution that contains some elongation activity-see lane 2) were added to immobilized templates and incubated for 15’, followed by a chase with unlabeled rNTPs. rP-TEFb plus a P11 elongation fraction (P.3c) was added to immobilized templates and compared to either no additional factors (lane 1) or the P.3c fraction alone (lane 2) as indicated in panel A. Transcription products were analyzed on an 8% urea/TBE gel followed by autoradiography. D. Purification of F-SEC. Flag-tagged SEC subunit AFF1 was purified from 293 cells using M2-agarose and eluted by excess Flag peptide. Input extract contained undetectable levels of SEC subunits, but which were concentrated by affinity purification (E1-E3), as indicated by western blot analysis for SEC subunits AFF1, ELL2, and CDK9. E. Kinase activity of F-SEC was assessed by incubating purified F-SEC (lane 2) or rP-TEFb (positive control, lane 3) with the known substrate GST-CTD, containing the first 26 repeats of human pol II CTD. Phosphorylation was detected by western blot using an anti-phosphoserine 2 CTD antibody. Lane 1 contains only the GST-CTD and the kinase reaction buffer. F. Schematic shows the steps in the immobilized template assay in panel G. G. F-SEC stimulates release of paused pol II. Either purified F-SEC or the control elution buffer were titrated into pulsed immobilized templates after they had been isolated and washed to remove ^32^P-CTP /GUA and nuclear extract. After a 15’ incubation, unlabeled NTPs were added for 5’ to chase any elongation competent pol II into making full-length RNAs. Transcription products were analyzed on an 8% urea/TBE gel followed by autoradiography. Vertical black line between lanes 1 and 2 indicates a deletion of intervening lanes relative to the original autoradiograph.

The Super Elongation Complex (SEC) physically connects the ELL elongation factor to the P-TEFb kinase (Luo et al. 2012; Lin et al. 2010). Both ELL and P-TEFb can stimulate elongation in vitro by themselves and the SEC can stimulate elongation as a discrete complex (Biswas et al. 2011). The question remains however, whether the SEC can stimulate pause release. Flag-tagged SEC (Lin et al. 2010) (F-SEC) was purified from 293 cells and western blot analysis showed that it contained several SEC subunits (E1-E3, Fig. 3D). The F-SEC phosphorylated rGST-CTD, as did rP-TEFb, as expected (Fig. 3E) (Biswas et al. 2011; Lin et al. 2010). Next, F-SEC was assayed using immobilized templates (Fig. 3F). Titration of F-SEC relative to the elution buffer (containing FLAG peptide) showed stimulation of isolated paused pol II complexes into intermediate and full-length RNA products (compare lanes 2-5 with 6-9, Fig. 3G). Thus, these stably isolated, engaged, paused polymerases can enter productive elongation with either P-TEFb or SEC.

These experiments all show that the CFS described here behaves as one would demand of a system to study RNA pol II pausing and elongation and to address the necessary factors and mechanisms of regulation. We now use this system to discover and explore previously unidentified factors in pausing and pause release.

### The OGT substrate UDP-GlcNAc blocks elongation in the CFS

Previously it was shown that OGT is required for PIC formation (Ranuncolo et al. 2012; Lewis et al. 2016) and that OGA is required for proper elongation in the CFS (Resto et al. 2016). Since OGA activity requires an O-GlcNAcylated substrate, prior OGT catalytic activity is also implicated in the transition to elongation.

To establish a direct requirement for O-GlcNAcylation in the CFS the substrate of OGT, UDP-GlcNAc, was titrated into the CFS (Fig. 4A). Titration of UDP-GlcNAc (after PIC formation) resulted in a concentration-dependent decrease in full-length RNA (lanes 1-4, Fig. 4A). A similar result was obtained with immobilized templates and the 0.3M P11 elongation fraction P.3c: addition of rOGT and UDP-GlcNAc decreased synthesis of full-length RNA (compare lanes 2 and 3 in Fig. 4B). That elongation was blocked suggested that pausing had increased via OGT activity and the O-GlcNAcylation of pausing factors.

**Figure 4.**
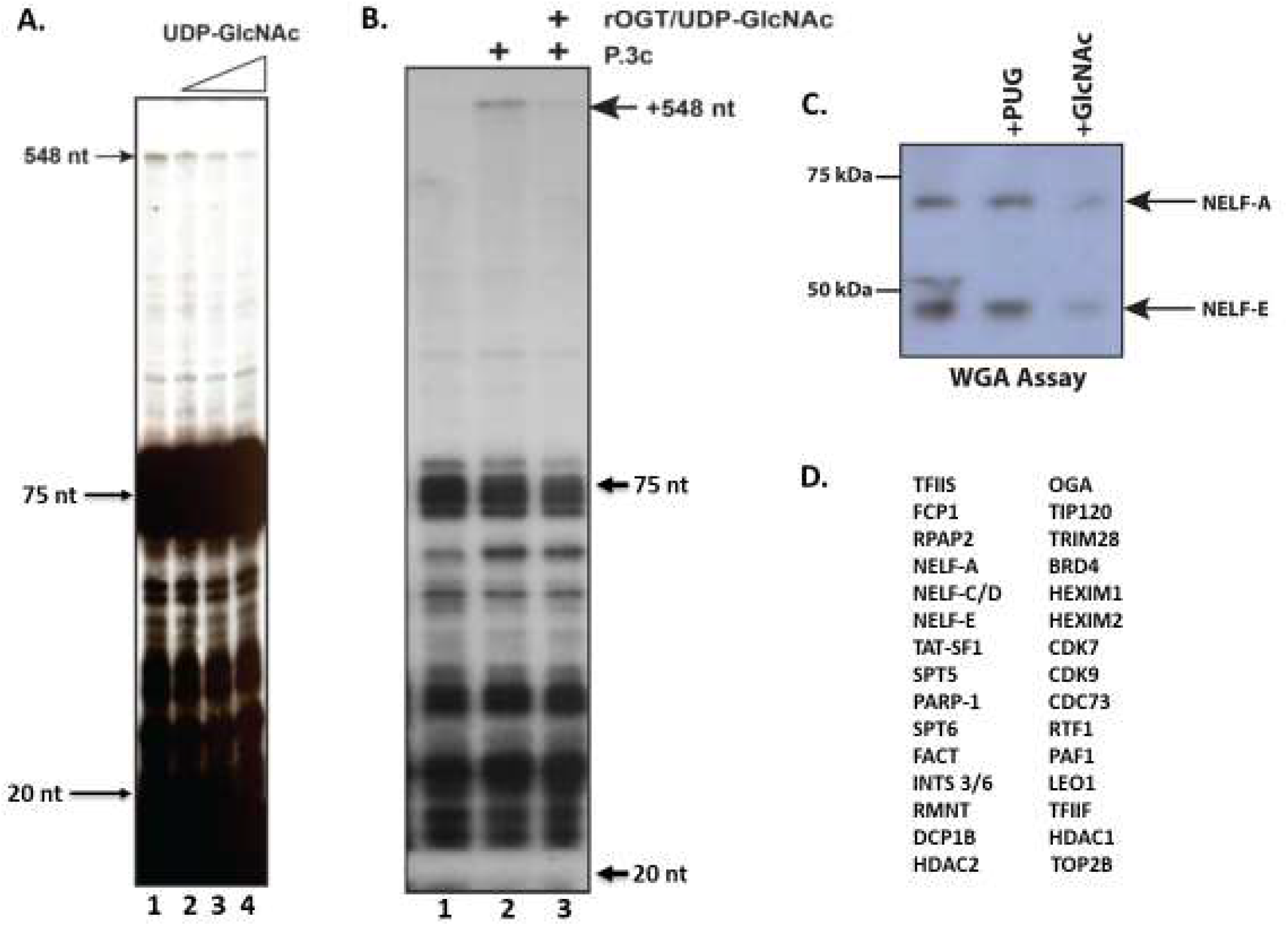
O-GlcNAcylation activity blocks RNA pol II elongation. A. UDP-GlcNAc causes a decrease in elongation in nuclear extracts. Standard pulse-chase assays were performed after incubating preformed PICs with increasing amounts of the OGT substrate UDP-GlcNAc (0.1, 0.2, 0.5, and 1 mM final concentrations). B. UDP-GlcNAc decreases elongation of EEC templates. Immobilized templates and pulsed EECs were isolated and washed in low salt transcription buffer. To the indicated reactions, either P.3c or P.3c plus 0.25 μg rOGT and 0.4 mM UDP-GlcNAc were added for 15’ prior to chasing labeled RNAs for 5’ with unlabeled NTPs. C. Pausing factor NELF is O-GlcNAcylated. Nuclear extracts (incubated as indicated with either 3 mM PUGNAc or 0.5M GlcNAc) were incubated with wheat germ agglutinin-agarose for 1 hour and washed several times. The recovered agarose beads were eluted in sample buffer, separated by PAGE, and transferred to nitrocellulose. NELF subunits were detected by western blot. D. Shown is a list of O-GlcNAcylated pausing and elongation factors that have been identified by mass spectroscopy(Hahne et al. 2012, 2013).

### Pausing and elongation factors are O-GlcNAcylated

To test if pausing factors are O-GlcNAcylated, nuclear extract was incubated with wheat germ agglutinin (WGA)-agarose beads to affinity purify O-GlcNAcylated proteins and the bound proteins were assayed by western blot. NELF-A and -E bound the WGA-beads (left-most lane, Fig. 4C) but did not bind beads that had been preincubated with the competitor GlcNAc, indicating that the WGA interaction was specific to O-GlcNAcylation (+GlcNAc, Fig. 4C). As a control for the possible loss of O-GlcNAcylation by endogenous OGA, the OGA inhibitor PUGNAc (+PUG; Fig. 4C) was added but did not alter the presence of O-GlcNAcylated NELF subunits. The literature for mass spectroscopy data of O-GlcNAcylated proteins further shows that at least 30 pausing and elongation factors are O-GlcNAcylated (Fig. 4D), including DSIF and NELF (Hahne et al. 2012, 2013).

### OGT inhibitors reduce pausing and allow RNA pol II to elongate

Previously, addition of the OGT catalytic inhibitor ST045849 (Gross et al. 2005) affected transcription (Ranuncolo et al. 2012), but at what step was not determined. Thus, we set out to examine the effects of OGT inhibition in a pulse-chase assay and to determine where in the transcription process OGT acted.

The addition of ST04 along with CMV promoter DNA at the start of PIC formation completely abrogated all RNA synthesis (compare lanes 1 and 2, Fig. 5A). However, ST04 added after PIC formation showed a release of pol II from the paused population into elongation (post-PIC, compare lanes 1 and 4, Fig. 5A), while PUGNAc showed the expected elongation defect (Resto et al. 2016) (lane 5, Fig. 5A). Titration of ST04 and a second OGT inhibitor ST06 showed similar effects: 0. 1mM ST04 and to a lesser extent 1mM ST06 showed that the paused pol II population at +20/+75 was released into longer RNAs (compare lane 2 to lanes 3 and 6, Fig. 5B).

**Figure 5.**
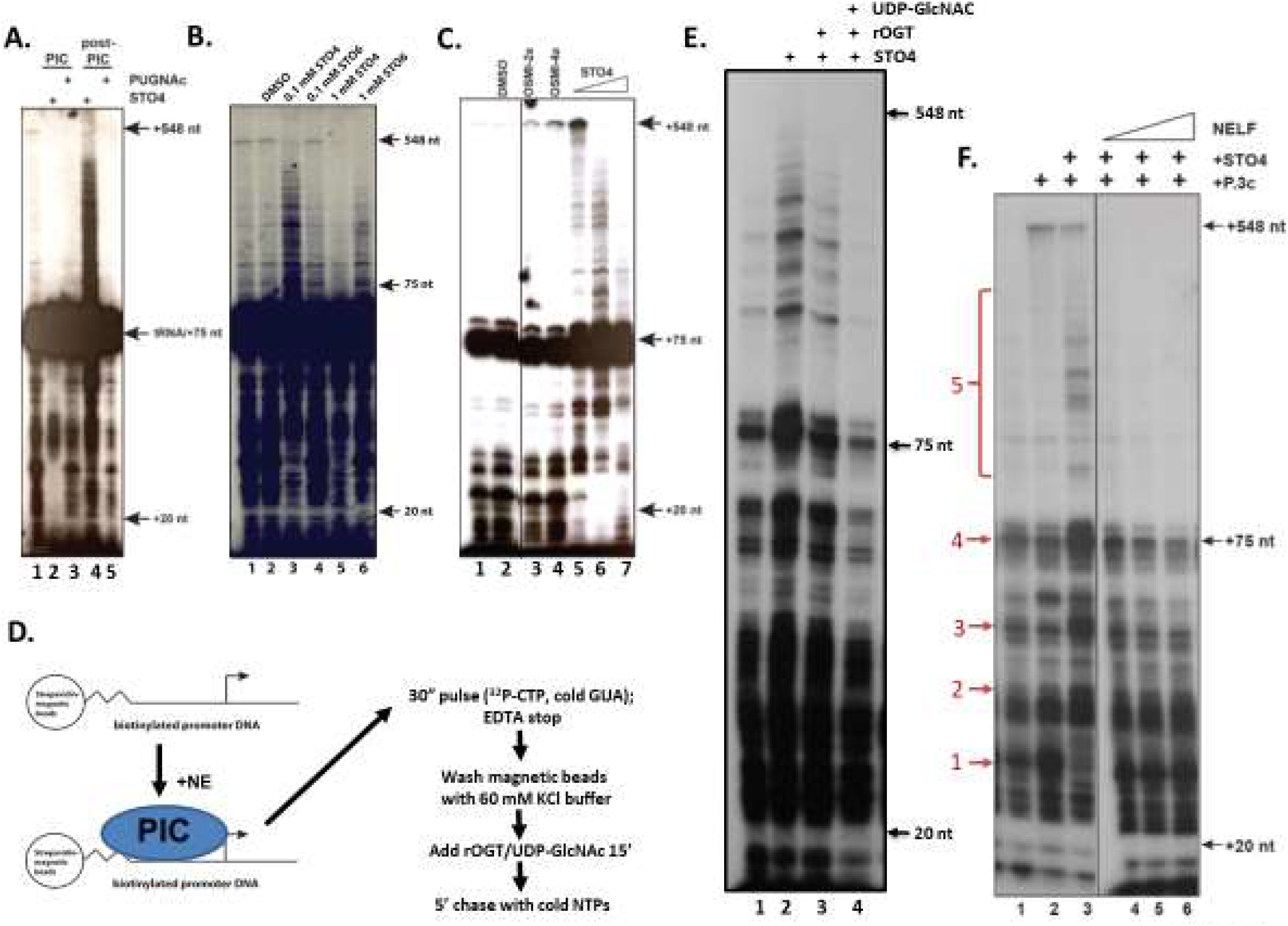
OGT is required to establish a paused pol II. A. OGT inhibitor addition after PIC formation causes release of paused pol II. 40 μM ST04 or 3mM PUGNAc were added simultaneously with CMV promoter DNA or 30’ afterwards (to allow PICs to form). These reactions were then assayed by standard pulse-chase assay. Transcription products were analyzed on an 8% urea/TBE gel followed by autoradiography. B. OGT inhibition causes release of paused pol II. The indicated final concentrations of either ST04 or ST06 were added to preformed PICs which were then assayed by pulse-chase. Transcription products were analyzed on an 8% urea/TBE gel followed by autoradiography. C. Three OGT inhibitors affect pause release. Either DMSO, 80 μM OSMI-2a or 160 μM OSMI-4a were added after PIC formation followed by a standard pulse-chase assay (lanes 1-4). Lanes 5-7 show a titration of STO4 (40, 80, 200 μM final concentrations). Transcription products were analyzed on an 8% urea/TBE gel followed by autoradiography. Vertical black line between lanes 2 and 3 indicates a deletion of intervening lanes relative to the original autoradiograph. D. Schematic shows the steps in the immobilized template assay in panel E. E. rOGT rescues the OGT inhibitor ST04 aberrant elongation effect. Immobilized templates were created by allowing PICs for form for 30’, followed by a 15’ incubation with 0.1 mM ST04 and then a 30’’ pulse with ^32^P-CTP /GUA followed by addition of EDTA to stop transcription. Beads were washed as indicated to remove all rNTPs, ST04, and nuclear extract. Individual reactions were suspended in transcription buffer and either rOGT or rOGT and 0.4 mM UDP-GlcNAc were added as indicated, for 15’, followed by a 5’ chase with unlabeled NTPs. Transcription products were analyzed on an 8% urea/TBE gel followed by autoradiography. F. NELF complex reverses the ST04-induced pause release. Immobilized template assays were set up similarly to panels D and E, but with the addition of the P.3c elongation fraction ± ST04 after the isolation of the pulsed EEC template, followed by the titration of F-NELF (lanes 4-6). After a 15’ incubation period, cold NTP were added as the chase step. Transcription products were analyzed on an 8% urea/TBE gel followed by autoradiography. Vertical black line between lanes 3 and 4 indicates a deletion of intervening lanes relative to the original autoradiograph.

More recently, the Walker lab synthesized several additional OGT inhibitors (Ortiz-Meoz et al. 2015). Titration of these inhibitors (OSMI-2a and OSMI-4a) post-PIC formation showed an increase in full-length RNA relative to the untreated and DMSO-treated nuclear extract (compare lanes 1 and 2 to lanes 3 and 4, Fig. 5C). Further refinement of the ST04 titration showed the same effects: escape of the paused pol II population, especially the population just above +20, and a significant increase in the full-length RNA product (lanes 5-7, Fig. 5C). As the ST04 concentration increased further, a second effect appeared closer to the full-length RNA at 548 nt (as seen in lane 4, Fig. 5A and lanes 3 and 6, Fig. 5B), suggesting a second OGT requirement for elongation into a full-length RNA at 548nt. Thus, four different OGT inhibitors show that OGT activity is necessary to establish a paused pol II, and in the absence of OGT, the pol II does not pause, but immediately moves into an elongation phase.

### OGT reverses the pausing defect of OGT inhibitors

If OGT is required for pausing, then the addition of rOGT and UDP-GlcNAc should reverse the effects of ST04. This experiment is readily done with immobilized templates. PICs were formed for 30’, then, where indicated, ST04 was added for 10’. Transcription was initiated with a 30” pulse followed by inhibition with 20 mM EDTA. The beads were then isolated using a magnet, washed to remove STO4, NTPs, and nuclear extract, and then resuspended in transcription buffer. To the indicated reactions either rOGT or rOGT plus UDP-GlcNAc were added for 15’, followed by a 5’ chase with cold NTPs (Fig. 5D). As expected, the untreated templates did not appreciably elongate after washing and the ST04 reaction showed much higher levels of RNAs longer than +75 (lanes 1 and 2, Fig. 5E). The addition of rOGT has a slight effect on the levels of these RNAs (lane 3, Fig. 5E) but the addition of both rOGT and its nucleotide-sugar substrate UDP-GlcNAc dramatically suppressed the levels of RNAs longer than +75 (lane 4, Fig. 5E), indicating that OGT catalytic activity reversed the pause-neutralizing effect of ST04. This experiment shows that OGT catalytic activity is directly necessary for establishment of the paused pol II.

### NELF rescues the pausing defect of OGT inhibition

The release of the paused pol II after treatment with ST04 suggested that the NELF pausing factor was specifically lost with OGT inactivation. To determine whether the addition of NELF back to a ST04-treated EEC could prevent the release of the paused polymerases, we again isolated paused pol II on immobilized templates (as in Fig. 5D), but after washing, added the P.3c elongation fraction with or without ST04 (lanes 1-3, Fig. 5F). With STO4 added, there was a clear release of paused pol II as indicated by the chasing of label from RNAs that are slightly larger than +20 (red arrow #1) into RNAs throughout the paused region (red arrows #2-4) and then into and beyond +75 (red bracket, #5; compare lanes 2 and 3, Fig. 5F). The titration of purified Flag-tagged NELF completely abrogated the release of these paused polymerases as indicated by the loss of all labeled RNA longer than +75 (red bracket, #5), as well as the reduction in RNA levels at and just below the +75 region (red arrows #2-4), and finally the retention of the short RNA population just above the +20 mark (red arrow #1) that had released into longer RNAs with STO4 (compare the RNA intensity indicated by arrow #1 between lanes 3 and 4-6). Mechanistically, these data suggest that O-GlcNAcylation is required for pausing by maintaining the paused pol II in a state receptive to NELF. Loss of O-GlcNAcylation causes the loss of NELF and release of the paused pol II into an elongation-competent species.

### The OGA inhibitor PUGNAc blocks elongation in vivo by suppressing RNA pol II pause release

To analyze the perturbation of O-GlcNAcylation on pausing and elongation in vivo, we used a variation on the SLAM-seq/TT-seq (Muhar et al. 2018; Schwalb et al. 2016). The incorporation of 4-thiouridine in vivo offered a rapid and efficacious approach to measuring changes in nascent RNA expression. The 4sU approach (Fig. 6A) requires only the isolation of the RNA after treatment of the cells with 4sU, followed by a brief treatment with iodoacetamide to alkylate the 4sU. During library PCR amplification, the alkylated 4sU is recognized as a C and thus a G is incorporated opposite the alkyl-4sU. During bioinformatic analysis the T > C conversions can be measured and isolated, identifying the pulsed, actively transcribed RNA population (Muhar et al. 2018). A library made with random primers permits the specific identification of nascent RNAs.

**Figure 6.**
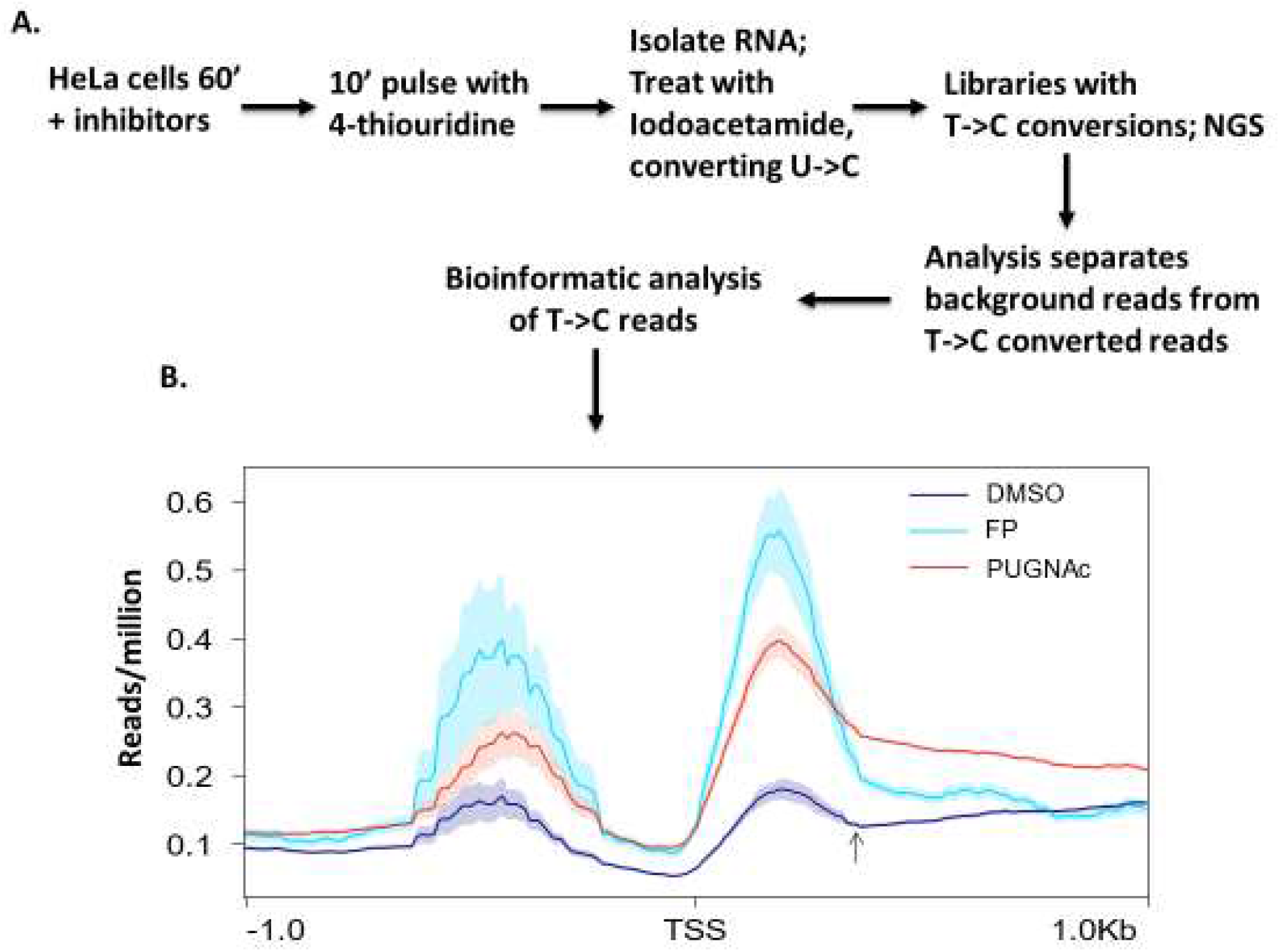
Inhibition of the O-GlcNAc aminidase blocks elongation in vivo. A. Schematic of the 4-thiouridine experiment. B. Metagene analysis of 4sU-incorporated RNA, mapping 4sU-RNAs based on distributions relative to the transcriptional start site (TSS) of genes, normalized to reads per million.

We treated HeLa cells with either the CDK9 inhibitor flavopiridol (FP) or the OGA inhibitor PUGNAc using concentrations known to have effects in vivo. As a negative control, HeLa cells were treated with an equivalent amount of DMSO. We used FP as a positive control as others have noted a significant reduction in elongation (Zhou et al. 2012) and an increase in the nascent RNA-defined paused pol II population at the 5’ end of genes after FP treatment (Laitem et al. 2015). As expected, metagene analysis of replicate FP treatments showed an increase in the paused pol II population just downstream of the transcriptional start site (TSS) followed by decreasing incorporation of 4sU, indicative of an elongation block (Figure 6B). Importantly, we found that the PUGNAc treatment resulted in a similar effect where pol II had accumulated above DMSO levels at the 5’ end of genes and a decreasing incorporation of 4sU. In contrast, the control DMSO-treated cells showed less accumulation of a paused pol II population and more importantly an upward sloping/increase in 4sU incorporation. These results show that OGA inhibition in vivo blocks elongation and promotes the accumulation of paused pol II at the 5’ end of genes, much like the behavior of flavopiridol, a known elongation inhibitor. Lastly, the requirement for OGA both in vivo and in the CFS requires, by definition, prior O-GlcNAcylation by OGT, and which we have shown to occur prior to OGA activity in the CFS (Fig. 5).

## Discussion

### Summary of the evidence of the recapitulation of RNA pol II pausing and elongation

We present here evidence that a human cell-free system (CFS) recapitulates the expected behaviors of pol II pausing and elongation. The CFS establishes a bona fide paused pol II, based on 12 different criteria (Figures 1-3). 1) The CFS synthesizes a population of paused RNAs found between +20 and +75 downstream of the TSS. 2) These RNAs are stably engaged to the DNA template and remain associated with the pol II. 3) The +20/+75 RNA population is intrinsically stable and alone will not chase into longer RNAs. 4) Sarkosyl treatment of this population on washed immobilized templates results in the release of the pol II and the synthesis of longer RNAs. 5) The treatment of the CFS with the CDK9 inhibitor flavopiridol inhibits elongation after +75 but does not perturb the +20/+75 RNA population. 6-7) The CFS is also dependent on CDK12/13 and PARP-1 catalytic activities, whose requirements were shown in vivo (Gibson et al. 2016; Zhang et al. 2016), but which remained to be seen in a direct, biochemical transcription assay. 8) In contrast, treatment with the CDK7 inhibitor, THZ1, eliminates the +20/+75 RNA population, as observed in other experiments (Nilson et al. 2015). 9) Excess SPT5 results in the synthesis of RNAs longer than +75, as expected if titrating out associated pausing factors. 10) The +20/+75 RNA population is released upon treatment with either rP-TEFb or purified SEC. 11) Using the CFS, we established paused pol II on the human heat shock promoter of the HSPA1B gene. 12) Price and colleagues recently showed that in vivo the CMV IE promoter used here and all other CMV promoters are in fact paused (Parida et al. 2019). These data show that the CFS establishes an engaged, paused RNA pol II population between +20 and +75 and which can enter productive elongation. Furthermore, we show that the human HSPA1B promoter, which harbors a paused pol II in vivo, also assembled a paused pol II in the CFS (Fig. 2).

These data make clear that chromatin and nucleosomes are not required for pausing per se. No chromatin has been added to the CFS nor have steps been taken to assemble any of these templates with nucleosomes. Our data also reflect CMV transcription in vivo, where pausing occurs at all viral promoters despite not being packaged into chromatin (Parida et al., 2019). Pausing is a direct, default consequence of transcription initiation.

### O-GlcNAc cycling is necessary for RNA pol II pausing and elongation

O-GlcNAc and OGA ChIP-seq peaks overlap paused pol II positions in human BJAB cells (Resto et al. 2016). OGT ChIP-seq peaks map to promoters in mice (Deplus et al. 2013) and OGT shRNA reduced pol II and O-GlcNAc levels at several human B-cell promoters (Ranuncolo et al. 2012). Mass spectroscopy data indicated that over 20 pausing and elongation-associated factors, including DSIF and NELF (Fig. 4), are O-GlcNAcylated (Hahne et al. 2012), as are the bovine and human RNA pol II C-terminal domains (Kelly et al. 1993; Ranuncolo et al. 2012). The modification of these factors accounts for the co-localization of O-GlcNAc and paused pol II at promoters in vivo. Previous experiments showed that inhibition of OGA in the CFS blocked elongation as effectively as the CDK9 inhibitor flavopiridol (Resto et al. 2016). Those experiments indicated that OGA catalytic activity was required for elongation and secondly, that OGT must have acted prior to that point in order to create a substrate for the OGA. The co-localization of OGT, OGA, and the O-GlcNAc modification at promoters in vivo, the reduction in pol II promoter occupancy after OGT knockdown, and the OGA requirement for elongation all point to a model where O-GlcNAc cycling regulates RNA pol II recruitment and elongation.

However, it was unclear at what steps in transcription OGT acted. The experiments in Fig. 5 precisely delineated OGT activity in both PIC formation and pause establishment. The rescue of the OGT inhibition with rOGT showed that pausing was a direct consequence of OGT catalytic activity. OGT inhibition was also rescued by the addition of purified NELF, strongly suggesting that the mechanism by which OGT established a paused pol II is via recruitment of NELF (Fig. 5). Furthermore, the titration of UDP-GlcNAc into the CFS blocked synthesis of a full-length RNA product, showing that increased O-GlcNAcylation decreased the elongation capacity of the CFS (Fig. 4). Likewise, the addition of rOGT and UDP-GlcNAc to an immobilized template assay also blocked elongation (Fig. 4). Lastly, we have confirmed the OGA, and by extension, the OGT requirement for elongation by showing a significant accumulation of paused pol II in vivo after OGA inhibition (Fig. 6).

Our results suggest a model where O-GlcNAcylation addition and removal respectively regulate the establishment of a paused pol II and the subsequent release of the paused pol II into productive elongation. O-GlcNAcylation may regulate phosphorylation by blocking access of kinases to the necessary serine and threonine residues. OGA activity removes this block and permits phosphorylation to occur, leading to pol II escape into productive elongation. Alternatively, O-GlcNAcylation addition/removal is directly regulating the activities of the pausing and elongation protein machinery.

### A multi-factor model of RNA pol II pausing and release

These data can be further assembled into a multifactor model of pol II pause establishment and release. Pol II pausing is established by DSIF and NELF in conjunction with the catalytic action of CDK7 (Larochelle et al. 2012) and OGT. Release of the paused pol II into productive elongation is also a multifactor release step requiring P-TEFb (Guo and Price 2013), OGA (Resto et al. 2016), and PARP-1 (Gibson et al. 2016) (Fig. 7). Lastly, the substrates ATP, UDP-GlcNAc, and NAD+ are all high energy compounds. Hence, we hypothesize that these high energy donors may satisfy specific energy requirements for pausing and pause release in addition to the effect of post-translational modification itself.

**Figure 7.**
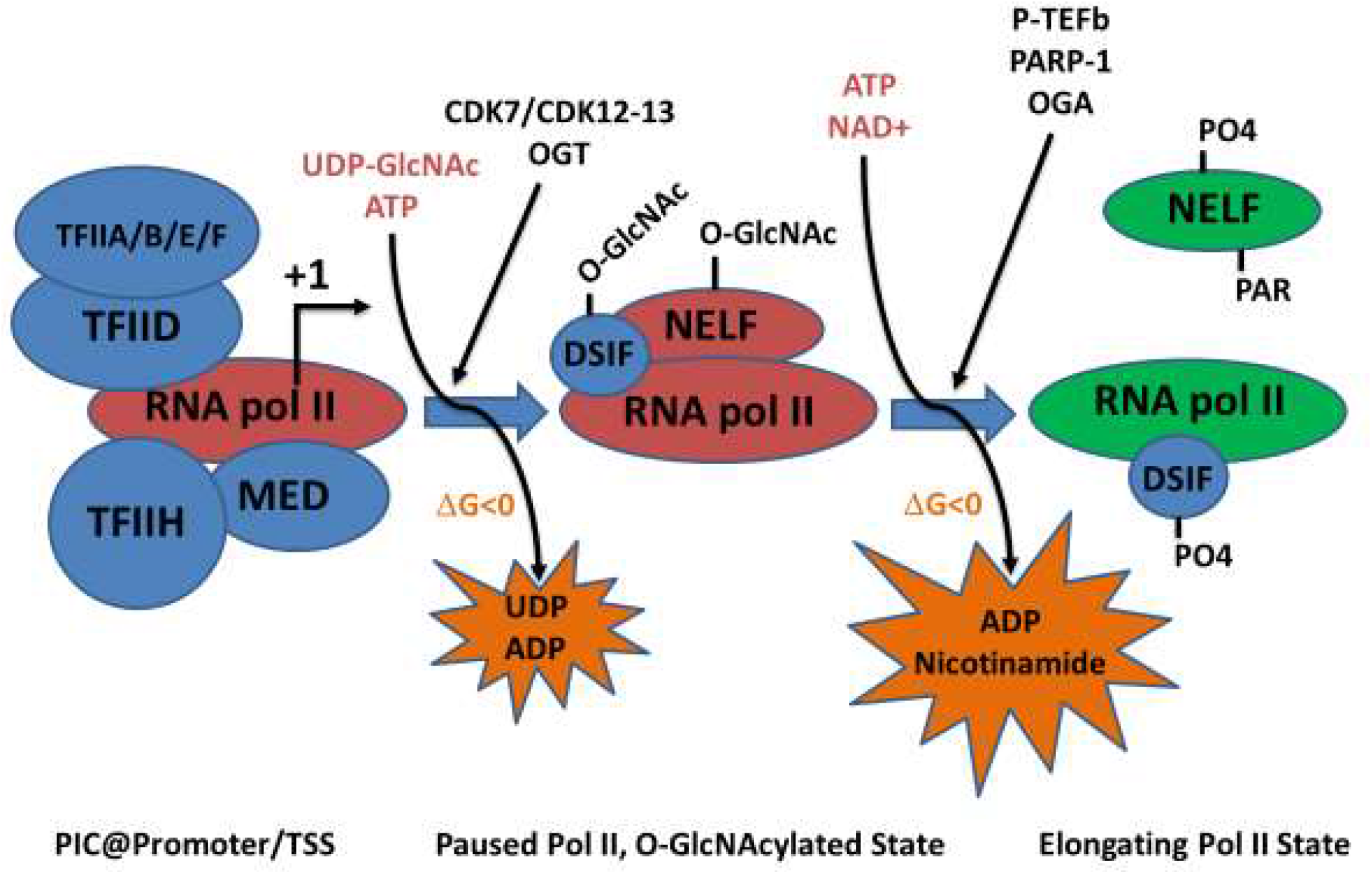
A multi-factor release model of RNA polymerase II pausing and elongation. The model proposes that there is a paused pol II complex in an O-GlcNAcylated state, partially defined by O-GlcNAcylated pausing factors DSIF and NELF and established by the concomitant catalytic activities of CDK7 (Larochelle et al. 2012), CDK12/13, and OGT. Release into productive elongation occurs by the action of P-TEFb (Guo and Price 2013), PARP-1 (Gibson et al. 2016), and OGA (Resto et al. 2016). This removes GlcNAc from the targeted proteins, poly-ADP ribosylates NELF (Gibson et al. 2016), possibly inducing NELF release and/or inactivation, and phosphorylates DSIF (Fujinaga et al. 2004; Ping and Rana 2001; Yamada et al. 2006; Peterlin and Price 2006), which converts DSIF into a positive elongation factor. The orange “bursting star” surrounding the various metabolites indicates that these hydrolysis products are associated with the release of free energy.

Furthermore, these high energy donor compounds are synthesized by various metabolic pathways. The OGT substrate UDP-GlcNAc is made from fructose-6-phosphate, glutamine, and acetyl CoA, derived from the glycolytic pathway, amino acid uptake, and lipid metabolism, respectively. The PARP substrate NAD+ is derived from tryptophan and connects oxidative phosphorylation and the Krebs cycle. Finally, ATP is the output from oxidative phosphorylation. If these enzymes are responsive to the metabolic flux of their substrates, then the pol II machinery at the 5’ proximal end of genes may be a sensor of the nutrient/metabolic state of the cell, modulating transcriptional output in response to changing metabolic flux.

### The cell-free transcription system

Pol II pausing is of paramount importance, as most genes now are known to be regulated at that stage and not at the earlier PIC formation/initiation stage. The lack of a human nuclear extract system that recapitulates pausing has hindered understanding of pausing. The cell-free system described here will allow further exploration of the depths of pol II pausing, the factors involved in this step, and how pausing and pause release might be regulated. Such a system allows one to directly probe these events, in contrast to the indirect genetic experiments to which one must resort for in vivo studies. Through the combination of the bottom-up functional biochemistry and in vivo genetic approaches, we should be able to further understand RNA pol II pausing and its regulation.

## Acknowledgments

The authors wish to thank Ali Shilatifard for the F-SEC cell line, Nathaniel Gray for THZ1 and THZ531, David Price for the AmSO_4_ extract, Yuki Yamaguchi for the GST-SPT5 expression vector and F-NELF HeLa cell line, Jane Jones of the NCI Frederick Protein Synthesis Core Facility for the production and isolation of rP-TEFb, Suzanne Walker for supplying the OSMI inhibitors, and the NCI Sequencing Facility for library construction and Illumina sequencing. B.A.L. wishes to especially thank Tobius Neuman for assistance in implementing the SLAM-DUNK bioinformatic analysis.

## Author contributions

B.A.L. conceived of the work, designed and conducted the experiments, analyzed the data, and wrote the manuscript. D.L. contributed to the data analysis and manuscript writing.

## Competing interests

The authors declare no competing interests.

### Data and materials availability

Raw and processed data are available in the GEO Repository, accession number GSE158746: https://www.ncbi.nlm.nih.gov/geo/query/acc.cgi?acc=GSE158746

## Materials and Methods

### In vitro transcription assays

Assays were all done (NE-based and EEC-based) as in the appropriate figure schematics and as described previously (Resto et al. 2016), except that a 60 mM KCl salt wash buffer (Cheng and Price 2007) was used to wash the EECs.

### Proteins, HeLa nuclear extract, and HeLa P.3 Fraction

rP-TEFb was synthesized in Tni-FNL cells by co-expression of CDK9-his6 and cyclinT1 subunits and then purified via IMAC. rGST and rGST-SPT5 proteins were purified as described previously for other GST-tagged proteins (Ranuncolo et al. 2012). HeLa nuclear extract was prepared as described, using a 0.4M KCl step for elution from nuclei to make the nuclear extract(Abmayr et al. 2006). F-SEC was purified from 293 Flp-in-TRex cells as indicated (Lin et al. 2010). rOGT and GST-CTD were purified as described previously (Ranuncolo et al. 2012). HeLa P.3 fraction was isolated by fractionation of a HeLa nuclear extract with a P11 column. The column was eluted with 0.1, 0.3, 0.5, and 1M KCl BC buffer (containing 20 mM Tris pH7.9 @ 4^°^C, 10% glycerol, 0.2 mM DTT, 0.2 mM EDTA). The 0.3M peak was pooled and precipitated with 50% ammonium sulfate. The resulting precipitate (P.3c) was suspended in 1/10^th^ volume (relative to the starting material volume) BC buffer without KCl and dialyzed against 0.1M KCl BC buffer. F-NELF complex was affinity-purified from stably transfected HeLa cells.

### WGA assay

20 μL wheat germ agglutinin (WGA) agarose beads was washed 1x with 500 μL BC100, spun to remove buffer with 26 gauge needle/syringe. 50 μL (10 mg/mL) nuclear extract was mixed with 50 μL H_2_O and incubated with the WGA beads for 60 minutes with rotation at room temperature. For the addition of PUGNAc, 10 μL of 20 mM PUGNAc was added to the 100 μL mixture of extract and H_2_O. To use GlcNAc as competitor, 50 μL 1M GlcNAc was added to the 50 μL extract and incubated with the WGA beads. After incubating for one hour, beads were washed 3x with BC100, eluted with Laemmli sample buffer, separated by denaturing PAGE, transferred to nitrocellulose, and analyzed for NELF by western blot.

### Kinase assay

0.2 μL GST CTD 1-26, 0.5 μL F-AFF1 or 0.5 μL rPTEFb, and 8.5 μL H_2_O, 1 μL 10x P-TEFb kinase buffer (50mM Tris pH7.5, 5mM DTT, 5mM MnCl_2_, 4mM MgCl_2_ (Zhou et al. 2000)) were mixed and incubated for 30 minutes at 30^0^C. Sample buffer was added to each sample and separated by denaturing PAGE followed by transfer to nitrocellulose and western blotting with anti-phosphoserine 2 antibody.

### 4sU experiment

Approximately, 1×10^7^ HeLa cells were grown on 10 cm plates and treated at 70% confluency with either DMSO or either flavopiridol (1 mM final) or PUGNAc (100 mM final) for 60 minutes. Cells were then treated with 1 mM final concentration of 4-thiouridine for 10 minutes. RNA was isolated using Trizol (Invitrogen) and treated with iodoacetamide as indicated (Herzog et al. 2017; Muhar et al. 2018). Libraries were constructed with indexed random primers and subjected to paired-end sequencing by the NCI Core Sequencing Facility. Two separate experiments were done using the same conditions. Sequences were analyzed by the SLAM-DUNK software (https://t-neumann.github.io/slamdunk/) (Neumann et al. 2019) and deeptools (https://deeptools.readthedocs.io/en/latest/index.html) (Ramírez et al. 2014). Raw and processed data are available in the GEO Repository, accession number GSE158746: https://www.ncbi.nlm.nih.gov/geo/query/acc.cgi?acc=GSE158746

### Antibodies

Anti-pol II CTD phosphoserine S2 (Abcam ab5095)

Anti-ELL2 (Abcam ab194445)

Anti-CDK9 (Bethyl A303-493A)

Anti-Flag (Sigma)

Anti-NELF-A (Santa Cruz sc-32911)

Anti-NELF-E (Santa Cruz sc-32912)

### Reagents

ST045849 and ST060266 (TimTec), Flavopiridol (Sigma F3055), PJ34, THZ531 (APExBIO, A8736), PUGNAc (Sigma A7229), ^32^P-CTP (3000 Ci/mmol, Perkin Elmer), NTPs (Roche), Sarkosyl (Sigma), WGA-agarose, GlcNAc (Sigma), UDP-GlcNAc (Sigma).

